# Laminar RNNs: using biologically-inspired network topology on the cortical laminar level in memory tasks

**DOI:** 10.1101/2024.05.06.592696

**Authors:** Ittai Shamir, Yaniv Assaf

## Abstract

Advancements in neuroscience and artificial intelligence have been fueling one another for decades. In this study, we integrate a neuroimaging model of laminar-level connectomics into a biologically-inspired deep learning model of recurrent neural networks (RNNs) for working memory tasks. The resulting model offers a way to incorporate a more comprehensive representation of brain topology into artificial intelligence without diminishing the performance of the network compared to previous models.

## Main

Neuroscience has propelled some of the earliest techniques in artificial intelligence (AI). The field of deep learning (DL) has drawn inspiration from some of the fundamental concepts of information processing in the human brain. The earliest DL network architectures consisted of simplified models of the brain, which included layers of neuron-like components (nodes) interconnected by weighted synapse-like connections (edges). One of the earliest DL networks was the multilayer perceptron (MLP), a straightforward DL model consisting of three layers: an input layer, a hidden layer, and an output layer (Murtagh 1991, LeCun et al. 2015). Since the MLP, the field of DL has expanded drastically to offer a wide variety of other discriminative (supervised) techniques, alongside a multitude of generative (unsupervised), hybrid learning, and other techniques. These techniques are based on different DL architectures, offering solutions for applications ranging from cybersecurity to healthcare (Sarker 2021). Nowadays, selecting and implementing a fitting DL architecture from the multitude of techniques available has become a challenge of its own. Adding to that challenge is the ‘black-box perception’ that often hinders the ability to interpret and understand these techniques (Sarker 2021). One approach for tackling these challenges has come once again from neuroscience, with renewed attempts to harness our increasing knowledge of the brain to develop artificial intelligence that better represents natural intelligence (Ullman 2019).

Over the last decade, great strides have been made in the field of neuroimaging with large-scale projects such as the Human Connectome Project (HCP), aimed at unraveling the architecture of the brain’s network of connections (Elam et al. 2021). Such projects have not only deepened our understanding of the brain, but also accelerated the mutually-beneficial relationship between neuroscience and DL, with advancements in both fields constantly fueling one another (Macpherson et al. 2021). On one hand, DL has fueled advancements in neuroscience with studies such as the one by Richie-Halford et al. (2022), which successfully used artificial neural networks for creating quality-controlled resources for connectomics research. On the other hand, neuroscience has fueled advancements in DL with studies such as the one by Szary et al. (2011), which developed biologically-inspired DL architecture for modeling general intelligence using reservoir computing.

Working memory is one of the central concepts in neuroscience and natural intelligence. Working memory is an executive cognitive function of the brain that relates to its capacity to track information over time (Slotnick et al. 2012). Recurrent neural networks (RNNs) are a popular deep neural network architecture that has an internal state designed for learning sequential or time-series data, which makes them particularly fitting for working memory tasks (Sarker 2021). RNNs, such as an Elman network, receive an input and continually feed the output of each previous step as input to the current one, creating a memory-like feature (Elman 1990, Vaswani et a. 2017).

In recent years, several open-source toolboxes have been created for building biologically-inspired RNNs and training and testing them on various tasks, including working memory tasks. These toolboxes include *conn2res*, which builds neuromorphic networks based on a derivation of RNNs called reservoir computing (Suárez et al. 2021, 2024), as well as *bio2art*, which builds bio-instantiated networks based on classic RNNs such as the Elmann network (Goulas et al. 2021, Damicelli et al. 2022). Concurrently, a new neuroimaging framework has introduced a model-based subfield of connectomics, termed ‘laminar connectomics’. The laminar connectome uses multimodal magnetic resonance imaging (MRI) and a data-driven model to derive a more nuanced characterization of the connectome that details connections on the level of cortical gray matter layers (Shamir and Assaf 2021a, 2021b, Shamir et al. 2022, Shamir and Assaf 2023). While biologically-inspired RNNs and laminar connectomics have vastly different goals, both are neuroimaging-based models that expand the knowledge of standard connectomics using both measured and assumed cortical connections. *bio2art* for example uses region-level connections from standard connectomics and assumes random (or selective) within-region connections (Goulas et al. 2021). The laminar connectome, on the other hand, models laminar-level connections based on multimodal neuroimaging techniques and assumes within-region connections based on previously published findings (Shamir and Assaf 2021a).

In this study, we integrate a multimodal neuroimaging model of laminar-level connectomics (Shamir and Assaf 2021b, Shamir et al. 2022,) into a biologically-inspired DL model of RNNs for various working memory tasks (Goulas et al. 2021, Damicelli et al. 2022) (see Fig. 1). Our hypothesis is that using laminar connectomes to integrate brain network topology on the cortical layer level in RNNs could offer a new level of detail and value to working memory tasks.

**Fig. 1:**
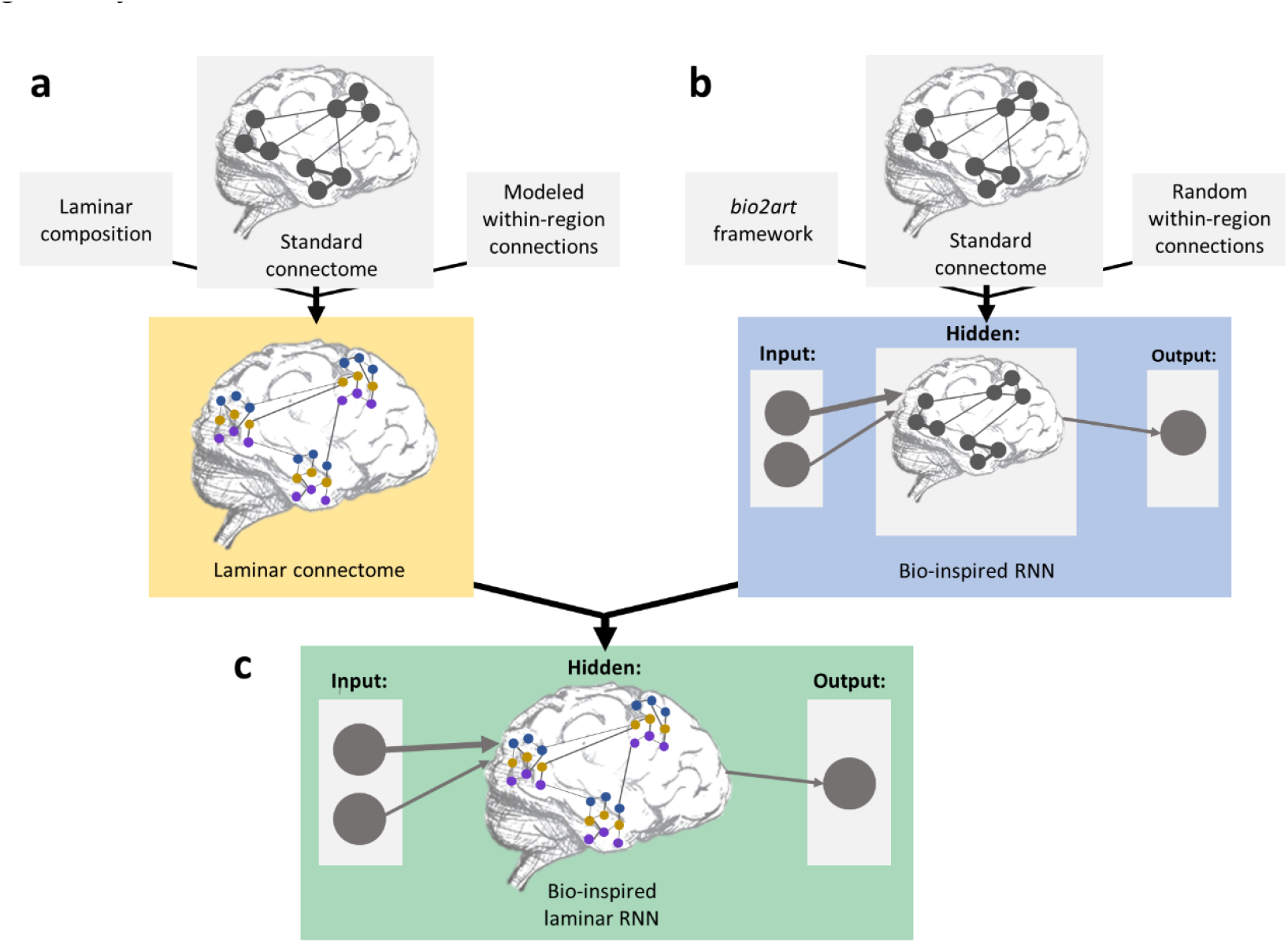
Integrating laminar connectomics into biologically-inspired RNNs: **(a)** Constructing a laminar connectome using multimodal neuroimaging datasets and a data-driven model, resulting in a laminar-level characterization of the connectome (Shamir and Assaf 2021a). **(b)** Constructing a biologically-inspired network using a standard connectomics neuroimaging dataset and the *bio2art* framework for converting biological networks into artificial RNNs (Goulas et al. 2021, Damicelli et al. 2022). **(c)** Constructing a biologically-inspired laminar-level RNN, using the laminar connectome in the *bio2art* framework.

To do so, we use an established approach for constructing bio-instantiated RNNs utilizing the computed standard and laminar connectomes of both human and macaque brains. The human connectomes represent the average connectivity patterns of (N=30) healthy humans, and the macaque connectome represents the connectivity patterns of an excised macaque brain (N=1). Datasets of standard and laminar connectomes for both species have been used and made public by Shamir and Assaf (2021b) (macaque dataset) and Shamir et al. (2022) (human dataset). We then use the *bio2art* framework to convert the connectomes into artificial RNNs, using the topology dictated by the biological neural networks (connectomes) and extrapolating from the empirical data (connection weights) to scale up for integration into an Elmann network (Elman 1990). The resulting RNNs are then trained and tested for various working memory tasks using the *bio_rnn_mem* framework (see Fig. 2). The working memory tasks involve predicting sequential data and include a sequence memory task and an N-back memory task. Both frameworks have been published and made available by Goulas et al. (2021) and Damicelli et al. (2022) (for more information, see the Methods section).

**Fig. 2:**
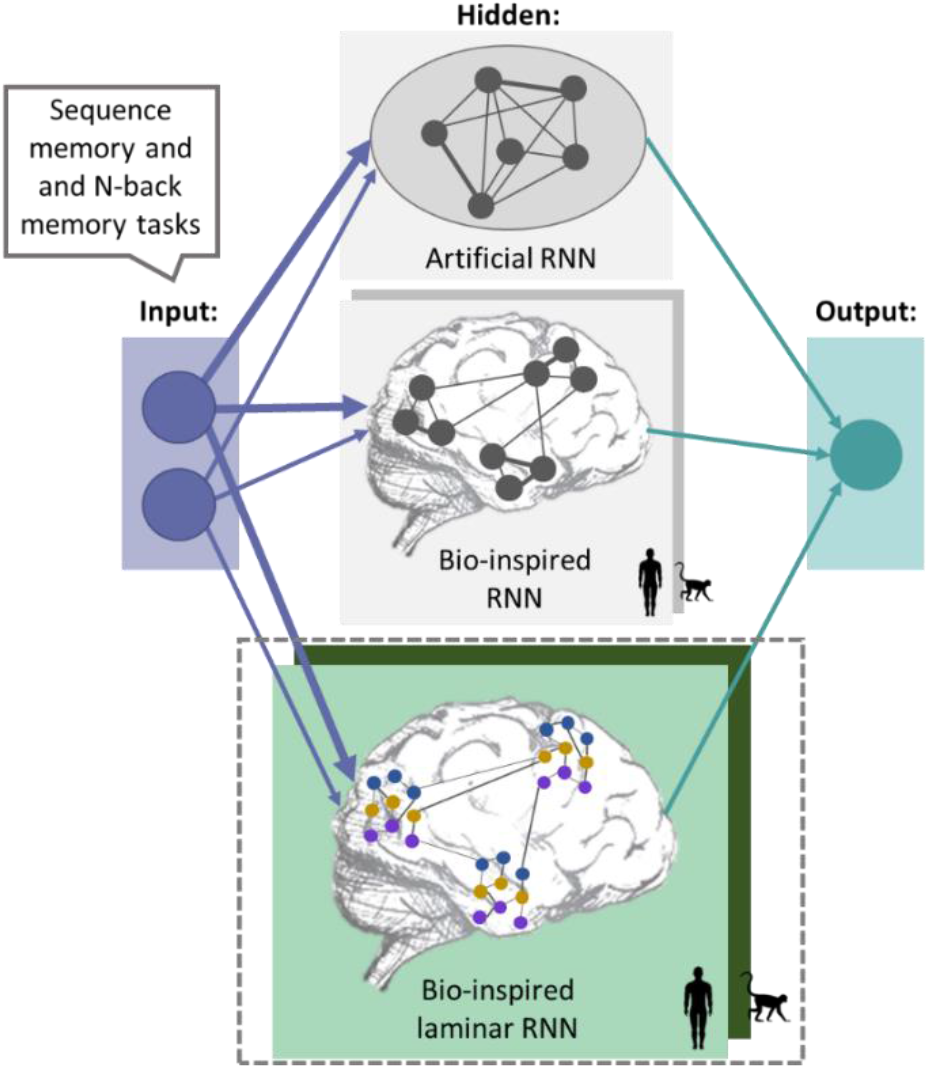
Exploring different types of RNNs for working memory tasks: Top: an artificial RNN based on random connectivity; middle: a biologically-inspired RNN using a whole-brain standard connectome, where each node represents a cortical region; bottom: a biologically-inspired RNN using a whole-brain laminar connectome, where each node represents a laminar component in a cortical region. The two biologically-inspired RNNs were constructed based on both human and macaque monkey connectomes. The RNNs were tested for working memory tasks, including a sequence memory task and an N-back memory task.

When constructing the different variations of biologically-inspired RNNs, experiments were conducted using a range of parameters, including learning rate, activation function, optimizer, and sequence length. Since the goal was to evaluate the prediction ability of RNNs based on different types of connectomes, these parameters were not optimized into a single hyperparameter combination. A similar choice was made by Goulas et al. (2021) and Damicelli et al. (2022), where they demonstrated that biologically-inspired RNNs based on standard connectomics can perform as well as artificial RNNs. To evaluate the potential benefit of using laminar-level connectomes, we chose to only focus on the following three network parameters:

Cortical atlas selection: The number of cortical regions in the atlas dictates the number of nodes in the network. In this study, we use atlases that parcellate the human cortex into 246 regions (Fan et al. 2016) and the macaque cortex into 167 regions (Felleman and Van Essen 1991), both based on connectional architecture. These atlases represent a finer parcellation compared to the ones used by Goulas et al. (2021), which include 57 regions (Betzel and Bassett 2018) and 29 regions (Markov et al. 2012), respectively.

1. Modeled within-region connections: We use the parameter N to represent the number of modeled neurons per cortical region. The previous study by Goulas et al. (2021) used N=4 neurons per region interconnected at random (set to 80% of all possible connections), after optimizing the RNN performance based on standard connectomics. In this study, we use N=1 for RNNs based on laminar connectomics, since within-region connections are built into the network based on the model of laminar connectivity. Each node in a laminar connectome represents a laminar component in a region, and so each cortical region is represented by three interconnected nodes. We use N=3 neurons per region (also interconnected at random) for RNNs based on standard connectomics, resulting in an equal total number of neurons (for comparability purposes).
2. Total number of neurons in the network: The total number of neurons in a network is the product of the number of regions and the number of within-region neurons. The selection of N=1 and N=3 for biologically-inspired RNNs based on laminar and standard connectomics (respectively), results in roughly an equal total number of neurons to previous studies, which assists in better comparing the performance of the RNNs.

The performance of the biologically-inspired laminar RNNs in both working memory tasks was then evaluated for both human and macaque monkey connectomes using loss and accuracy measurements (see Fig. 3). Initial examination of the results showed no considerable advantage to one cortical atlas over another, indicating no advantage to increasing the number of cortical regions. However, a slight advantage can be seen in including multiple neurons per cortical region over a single neuron per region (see Fig. 3a). In some cases, a slight advantage can be seen for laminar RNNs in loss minimization (see Fig. 3b), as well as in accuracy values (Fig. 3d). Loss and accuracy values of laminar RNNs seem to match those reached by RNNs based on standard connectomics (see Fig. 3c). Overall, results for laminar RNNs in both working memory tasks indicate no diminished performance compared to previous findings.

**Fig. 3:**
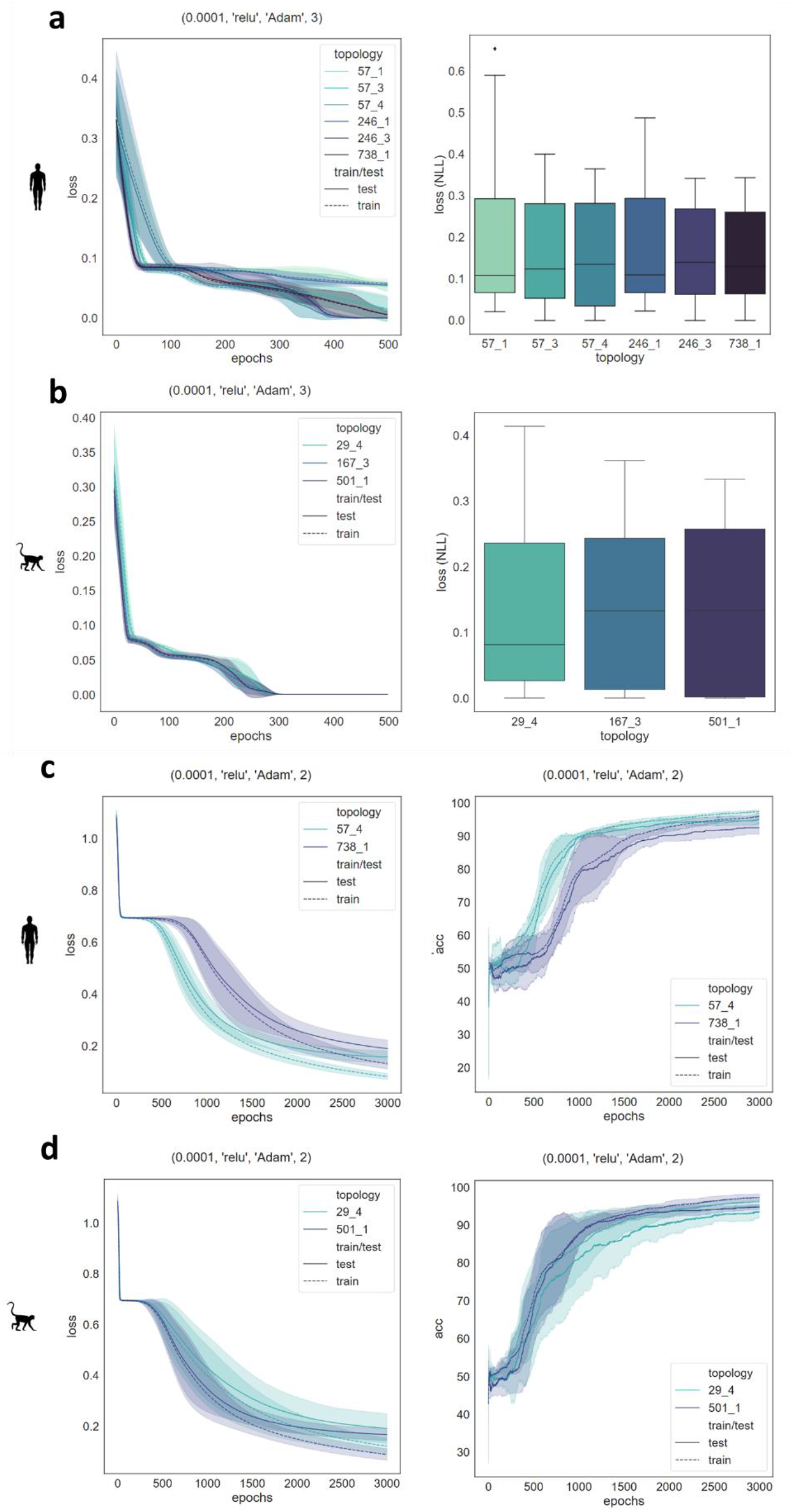
Evaluating the performance of standard and laminar RNNs: The RNNs examined are biologically-inspired and include those based on either standard or laminar connectomes. Each RNN examined is specified by two numbers: N1_N2, where N1 represents the number of atlas regions, and N2 represents the number of neurons per region. Human connectomes include 57 and 246 regions for standard connectomes, and 738 regions for the laminar connectome. Macaque connectomes include 29 and 167 regions for standard connectomes, and 501 for the laminar connectome. Results across epochs are shown for one parameter combination, denoted at the top by (learning rate, activation function, optimizer, sequence length). Shaded areas across epochs denote the standard deviations across five different runs of the experiment using the same parameters. Error bars denote the variability in results using a single connectome across twelve unique combinations of the above-mentioned parameters. (a) and (b) include results for a sequence memory task, including loss across epochs (left) and negative log likelihood (NLL) loss (right). **(a)** Sequence memory task results for RNNs based on various human connectomes. **(b)** Sequence task results for RNNs based on various macaque monkey connectomes. (c) and (d) include results for an N-back memory task, including loss across epochs (left) and accuracy across epochs (right). **(c)** N-back memory task for RNNs based on various human connectomes. **(d)** N-back memory task results for RNNs based on various macaque monkey connectomes.

The laminar structure of the cortex has long been assumed to play an important role not only in the development and various pathologies of the brain, but also its function in general and memory-related functions in particular (Shamir and Assaf 2023, Adesnik and Naka 2018). Various models have been proposed for illustrating the role of the cortical layers in memory functions. For example, various memory models of the somatosensory cortex assign layers either specific roles, such as sensory and motor memory storage (in layers II-III and V respectively), or computations, such as motivation, storage, and movement (Adesnik and Naka 2018).

In this study we integrate two intricate models from the fields of multimodal neuroimaging (Shamir and Assaf 2023) and biologically-inspired DL memory architectures (Goulas et al. 2021, Damicelli et al. 2022). The ensuing hybrid model is based on a comprehensive network model of brain connectivity on the cortical layer level. In this hybrid, aptly named a laminar RNN, between-region connections are modeled on the laminar level and within-region connections are modeled according to empirical findings. A new level of detail is introduced by using a laminar connectome, and simultaneously the complexity of the DL model is reduced by using a single neuron per region. Despite this apparent complexity reduction, using laminar RNNs indicates no diminished performance in working memory tasks compared to previous findings.

This study is a call to the neuroscience community to continue pushing the boundaries and integrating novel models from both neuroimaging and deep learning to develop better artificial intelligence and possibly gain a deeper understanding of natural intelligence. Integrating multi-faceted and multimodal models from both fields holds the potential to help bridge the gap between natural and artificial intelligence and benefit the scientific community at large.

## Methods

### Connectomes

#### Image acquisition

##### 1. Human connectome

An average connectome of N=30 subjects, 17 female, 18-78 y/o, neurologically and radiologically healthy (Shamir et al. 2022). The study was approved by the institutional review boards of Sheba Medical Center and Tel Aviv University. Subjects were scanned in a 3T Magnetom Siemens Prisma scanner with a 64-channel RF coil and gradient strength of up to 80 mT/m at 200 m/T/s:

a. A T1-weighted MPRAGE sequence: TR/TE = 1750/2.6 ms, TI = 900 ms, 1×1×1 mm^3^, 224×224×160 voxels.
b. An IR EPI sequence: TR/TE = 10,000/30 ms and 60 inversion times between 50 ms up to 3,000 ms, 3×3×3 mm^3^, 68×68×42 voxels, each voxel fitted with up to 7 T1 values.
c. A standard DWI sequence: Δ/δ=60/15.5 ms, b max=5000 (0 250 1000 3000 & 5000) s/mm^2^, with 87 gradient directions, FoV 204 mm, maxG= 7.2, TR=5200 ms, TE=118 ms, 1.5×1.5×1.5 mm^3^, 128×128×94 voxels. One B0 image was acquired with reversed phase encoding for EPI distortion correction.

##### 2. Macaque connectome

N=1 excised macaque brain obtained from the Mammalian MRI (MaMI) database (Shamir and Assaf 2021b). No animals were deliberately euthanized for the present study. The brain was scanned in a 7 T/30 Bruker scanner with a 660 mT/m gradient system:

a. A T1w sequence with a 3D modified driven equilibrium Fourier transform (MDEFT): TR/TE = 1300/2.9 ms, TI = 400 ms, 0.2×0.2×0.2 mm^3^, 300×360×220 voxels.
b. An IR sequence using 3D FLASH: TR/TE = 1300/4.672 ms and 44 inversion times between 25 ms up to 1,000 ms, 0.67×0.67×0.67 mm^3^, 96×96×68 voxels, each voxel fitted with up to 8 T1 values.
c. A DWI sequence with 4 segments diffusion weighted EPI sequence: 0.48×0.48×0.48 mm^3^, 128×160×116 voxels, Δ/δ = 20/3.3 ms, b = 5000 s/mm2, with 96 gradient directions and additional 4 with b=0.

For both datasets, the first two sequences were used for cortical laminar composition analysis, the third was used for mapping standard connectomics, and together they were used for modeling the laminar connectomes (Shamir and Assaf 2021a, 2021b, Shamir et al. 2022, Shamir and Assaf 2023).

#### Atlas selection

##### 1. Human atlas

The average human connectome was analyzed across the Brainnetome, a brain atlas based on connectional architecture, parcellated into 246 cortical regions (Fan et al. 2016). Subnetworks were analyzed based on an atlas of intrinsic functional connectivity, parcellated into seven subnetworks: visual, somatomotor, dorsal attention, ventral attention, limbic, frontoparietal, and default networks (Yeo et al. 2011).

##### 2. Macaque connectome

The macaque brain was analyzed across the FVE91 atlas, an atlas based on histological findings regarding connectional architecture, parcellated into 167 cortical regions (Felleman and Van Essen 1991).

#### Bio-instantiated RNN modeling

We used an established approach for constructing bio-instantiated RNNs (Goulas et al. 2021, Damicelli et al. 2022) utilizing the resulting laminar connectomes of both human and macaque brains (see figure 1). Code for creating (bio2art) and training these RNNs available at: https://github.com/AlGoulas/bio2art https://github.com/AlGoulas/bio_rnn_mem (respectively).

These studies use a previously published human standard connectome based on long-distance interareal connections, parcellated into 57 cortical regions (Betzel and Bassett 2018). The studies also use a previously published macaque standard connectome based on interareal connections found using histology, parcellated into 29 cortical regions (Markov et al. 2012).

#### Working memory tasks

The following working memory tasks were performed on various networks and subnetworks:

##### 1. Sequence memory task

A sequence of N numbers is generated uniformly and randomly and fed into the RNN in the memorize phase. Once a cue is provided, the network then tries to generate the same sequence of N numbers in the recall phase. The loss function is calculated according to the mean square error (MSE) between the predicted and the actual sequences.

##### 2. N-back memory task

A sequence of M numbers is generated uniformly and randomly and fed into the RNN in the memorize phase. Once a cue is provided, the network then tries to determine whether the last number is the same number N steps before. The loss function is calculated according to the negative log-likelihood between the correct response and the network output.

To determine the effect of using connectomes with different levels of detail, both working memory tasks were tested on whole-brain networks using both standard and laminar-level connectomes for both human and macaque monkey brains.

